# Strong Reward Signals, Weak Transfer: Limits of Spatial Priority Map Plasticity Across Task Contexts

**DOI:** 10.64898/2026.03.06.710060

**Authors:** Olgerta Asko, Maria Stavrinou, Thomas Hagen, Thomas Espeseth

## Abstract

Reward learning can bias attentional selection, but whether spatially biased reinforcement produces durable, context-general changes in spatial priority over days, and what neurophysiological signals track such learning, remains uncertain. We combined electroencephalography (EEG) and pupillometry with a multi-session spatial reward-learning paradigm (Chelazzi et al., 2014) in which targets could appear at eight locations and reward probability was systematically biased across locations during two days of training. A separate baseline/test visual detection task was administered before training and again four days after training to assess delayed transfer under cross-target competition. Training produced strong reward signals across measures. Feedback-locked ERPs (FRN and P300) differentiated outcome valence and reward magnitude and varied systematically with time-on-task, while pupil dilation was larger following high- than low-reward feedback and overall task-evoked responses decreased across blocks. Reward history also modulated stimulus-locked target processing: targets at high- versus low-reward locations differed reliably across N1/N2 and a late positivity, indicating multi-stage value-dependent influences on visual processing during active learning. In contrast, transfer was weak in both behavior and ERPs: behavioral indices did not show a reliable advantage for highly rewarded locations at delayed test, and neural evidence for persistence was limited to an N2 modulation in the most diagnostic high-versus-low comparison, which should be interpreted cautiously given low trial counts. Accordingly, we did not replicate the robust long-term spatial priority effect reported in the original study. Together, these findings reveal strong reward-learning signals but weak cross-task transfer, suggesting limits on how readily spatial reward learning consolidates into persistent, task-general spatial priority-map plasticity across contexts.

## Introduction

Motivational processes shape human behavior, learning, and skill acquisition. External rewards (e.g., food, money) can increase the motivational value of stimuli and actions and promote learning through reinforcement mechanisms (Berridge & Robinson, 2003; Dayan & Balleine, 2002; Schultz et al., 1997). Reward learning is also relevant for maladaptive patterns such as compulsive behaviors and addiction, where previously rewarded cues and contexts can bias attention and action beyond the rewarding episode (Field & Cox, 2008; Robinson & Berridge, 2008). These observations motivate research not only on *whether* reward biases selection, but on *how* such biases develop, consolidate over time, and generalize across contexts (Anderson, 2016; Failing & Theeuwes, 2018).

A useful contemporary framing is that attentional selection reflects the joint influence of current goals, physical salience, and selection history, lingering biases produced by prior experience, including reward learning. This tripartite perspective helps situate value-driven attentional biases alongside other learned regularities (e.g., probability cuing, statistical learning) and clarifies why reward effects can appear “automatic” even when no longer goal-relevant (Anderson et al., 2021; Awh et al., 2012; Theeuwes, 2019).

A substantial body of work has characterized short-term effects of reward on attentional selection: stimuli, features, or locations previously associated with reward can capture attention and gain priority even when they are no longer task-relevant (Anderson, 2013; Anderson et al., 2011; Della Libera & Chelazzi, 2009; Hickey et al., 2010; Libera & Chelazzi, 2006; Peck et al., 2009). At the same time, recent work highlights that the reliability of commonly used value-driven capture metrics can vary with task design, scoring choices, and analytic flexibility, an important consideration when interpreting individual differences or comparing effects across sessions and paradigms (Garre-Frutos et al., 2024; Stanković, 2025). Despite this progress, less is known about the temporal dynamics of reward-driven attentional biases beyond the immediate session, particularly their consolidation and persistence across days, and generalization across changes in task context and stimulus material.

Most evidence for value-driven attention has focused on feature-based biases, but reward can also bias selection in the spatial domain. Chelazzi et al. (2014) provided behavioral evidence that associating specific spatial locations with higher reward can produce long-lasting changes in spatial priority in a way that persists after rewards are discontinued and can generalize across changes in task and stimuli. These findings suggest that reward can shape spatial priority in a way that outlasts immediate reinforcement. Subsequent work has emphasized that the persistence and transfer of reward-history effects can be context-dependent: learned attentional biases may be strong within the trained environment or task structure yet attenuate or change when the broader context, stimulus set, or task demands shift (Anderson & Britton, 2019; Anderson & Kim, 2018; Bourgeois et al., 2018).

Spatial priority maps are typically conceptualized as topographic representations that integrate multiple biasing signals, such as stimulus-driven salience, current goals, and learned value, to guide selection. Stimulus-driven contributions are often linked to early visual and subcortical pathways (Itti & Koch, 2001), whereas value and control signals involve fronto-striatal and cingulo-opercular circuits supporting reward evaluation and action selection (Kennerley et al., 2011). The persistence of learned biases is thought to recruit memory- and reinforcement learning related systems, including medial temporal and striatal circuitry (Hutchinson & Turk-Browne, 2012; Pennartz et al., 2011). Contemporary accounts emphasize that learned value biases orienting by modulating priority signals within attention networks rather than solely altering overt choice (Itthipuripat et al., 2019). Together, these interactions support a distributed priority architecture in which reward history can shape the competition among locations and objects (Klink et al., 2014; Krauzlis et al., 2014).

Two issues remain underspecified. First, the durability and cross-context transfer of reward-biased spatial selection over multi-day intervals is not well established. Second, the time-resolved neurophysiological markers that accompany the formation of these biases, and potentially their later expression, are less well characterized, especially when behavior provides an ambiguous readout. These questions motivate the use of temporally precise measures that capture both reward processing and attentional selection.

Electroencephalography (EEG) offers a temporally precise window onto both outcome evaluation and stimulus selection. In outcome evaluation, event-related potentials (ERPs) such as the feedback-related negativity (FRN) and the P3 have been widely used to index how the brain registers feedback valence and reward magnitude over time (San Martín, 2012). Within reinforcement-learning accounts, the FRN has been linked to reward prediction error signals that influence medial frontal systems, including anterior cingulate cortex (ACC), supporting adaptive adjustments in behavior (Holroyd & Coles, 2002; Holroyd & Yeung, 2012). The P3 (including subcomponents often discussed as P3a/P3b depending on context) is commonly associated with updating and allocation of processing resources and is sensitive to motivational significance and outcome evaluation. Beyond feedback-locked components, stimulus-locked ERPs can index stages of target processing and selection, and reward history has been shown to modulate these responses. The N2pc has been widely used to index attentional selection in visual search, with reward history modulating N2pc amplitude for rewarded stimuli (Harris et al., 2016; Qi et al., 2013; Taylor & Feldmann-Wüstefeld, 2024; Wirth & Wentura, 2023). More generally, stimulus-locked modulation across early sensory, intermediate selection/control, and later updating stages can help localize *when* reward history influences processing.

The N2 component, particularly over frontocentral sites, has been linked to conflict monitoring and cognitive control processes (Botvinick et al., 2001; Folstein & Van Petten, 2008; Yeung et al., 2004). According to conflict monitoring theory, the ACC detects response conflict (i.e., the coactivation of mutually incompatible actions) and signals the need for increased attentional control (Botvinick et al., 2001; Kerns et al., 2004). This function may be particularly relevant for understanding how reward history influences selection when competing targets appear at differentially rewarded locations. An alternative account suggests that ACC activity reflects predictions of error likelihood based on reinforcement learning (Brown & Braver, 2005). Dissociating these accounts has proven challenging, but empirical evidence suggests that N2 amplitude varies inversely with error rate, consistent with conflict monitoring rather than error likelihood predictions (Nieuwenhuis et al., 2003; Yeung & Sanfey, 2004).

Pupillometry provides a complementary measure of reward-related processing and arousal/effort dynamics. Task-evoked pupil responses can scale with reward and track learning-related changes over time, but pupil diameter is also influenced by multiple cognitive and sensory factors. Accordingly, it is best interpreted as an integrative autonomic index rather than a one-to-one readout of a single neuromodulator (Cole et al., 2022; Megemont et al., 2022). The locus coeruleus-norepinephrine (LC-NE) system is thought to play a key role in modulating pupil diameter, with LC activity increasing neural gain and enhancing processing of task-relevant stimuli (Aston-Jones & Cohen, 2005; Joshi et al., 2016; Reimer et al., 2016). However, recent work has emphasized limits on one-to-one mapping between pupil diameter and LC spiking on a moment-by-moment basis, while still supporting meaningful links between arousal, learning variables, and pupil dynamics at appropriate timescales (Cole et al., 2022; Megemont et al., 2022). Combining EEG and pupillometry can therefore help dissociate outcome-evaluation dynamics from changes in alertness/effort across training.

In the present study, we adopted the multi-day spatial reward-learning paradigm introduced by Chelazzi et al. (2014), and extended it with concurrent EEG and pupillometry to characterize reward effects during active learning and to test for transfer under changed conditions. Across eight spatial locations, participants completed (i) a rewarded training task designed to bias spatial priority through differential reinforcement and (ii) separate baseline/test sessions administered days apart that assessed whether any learned spatial bias is detectable after a delay and across changes in task/stimulus context. During training we analyzed feedback-locked ERPs (FRN and P3) and pupil responses as a function of feedback valence, reward magnitude, and training progression, and we examined stimulus-locked ERPs for location-related modulations during target processing. During baseline/test we assessed whether any location-related ERP modulation is detectable after the delay in the separate task context.

Our aims were to (a) test whether reward training produces persistent spatial priority effects and whether such effects generalize to a related but distinct task/stimulus context after several days; (b) characterize feedback-related ERP signatures of outcome evaluation during training as a function of valence, magnitude, and time-on-task; (c) determine whether reward-associated locations modulate stimulus-locked ERPs during training and whether any modulation is detectable at delayed test/stimulus change; and (d) examine whether task-evoked pupil responses scale with reward magnitude and track learning dynamics across training.

We expected that reward learning would bias spatial processing such that targets appearing at highly rewarded locations would be processed more efficiently than targets at low-reward locations. This was expected to be reflected behaviorally and in stimulus-locked ERPs indexing target processing/selection, with differences between high- and low-reward locations. We also expected robust reward sensitivity in feedback-locked ERPs during training, with FRN and P3 modulated by feedback valence and reward magnitude and potentially varying with training progression, and convergent modulation in pupillometry. Because precise component-level predictions for spatial reward biases were not established, we did not have strong a priori predictions about which stimulus-locked components would be most sensitive and anticipated that reward history could modulate processing across early sensory, selection-related, and later evaluative or updating stages.

## Results

Forty healthy participants completed a four-session protocol: a baseline visual search task (Day 1), two days of reward-based training (800 trials/day; 1,600 total) with location-dependent reward probabilities (Days 2–3), and a delayed test session identical to baseline administered 4 days later (Day 7). The baseline/test task comprised 640 trials per session and assessed spatial target selection under single- and double-target conditions, including competitive trials intended to reveal reward-history biases. During training, reward feedback was delivered at the target location according to a fixed schedule that created high- and low-reward spatial positions. We recorded EEG throughout all sessions to quantify target-locked and feedback-locked ERPs, and recorded pupil diameter during training to index sensitivity to reward and time-on-task. Results are reported first for behavior, then for ERPs, and finally for pupil responses, distinguishing robust within-training learning signals from delayed transfer effects in the baseline/test context.

### Behavior

#### Training sessions

Behavioral analyses followed the approach of Chelazzi et al. (2014). During the two sessions of training (Days 2-3), target-report accuracy (ACC) increased from an average of 84.50 ± 1.27 % standard error of the mean (SEM) on the first training session to 91.68 ± 0.79 % on the second session, while reaction times (RTs) decreased from 707.74 ± 16.71 ms to 658.49 ± 14.21 ms. For visualization and analysis, each training day was divided into two blocks, yielding four blocks in total (400 trials per block; Figure 1A).

**Figure 1:**
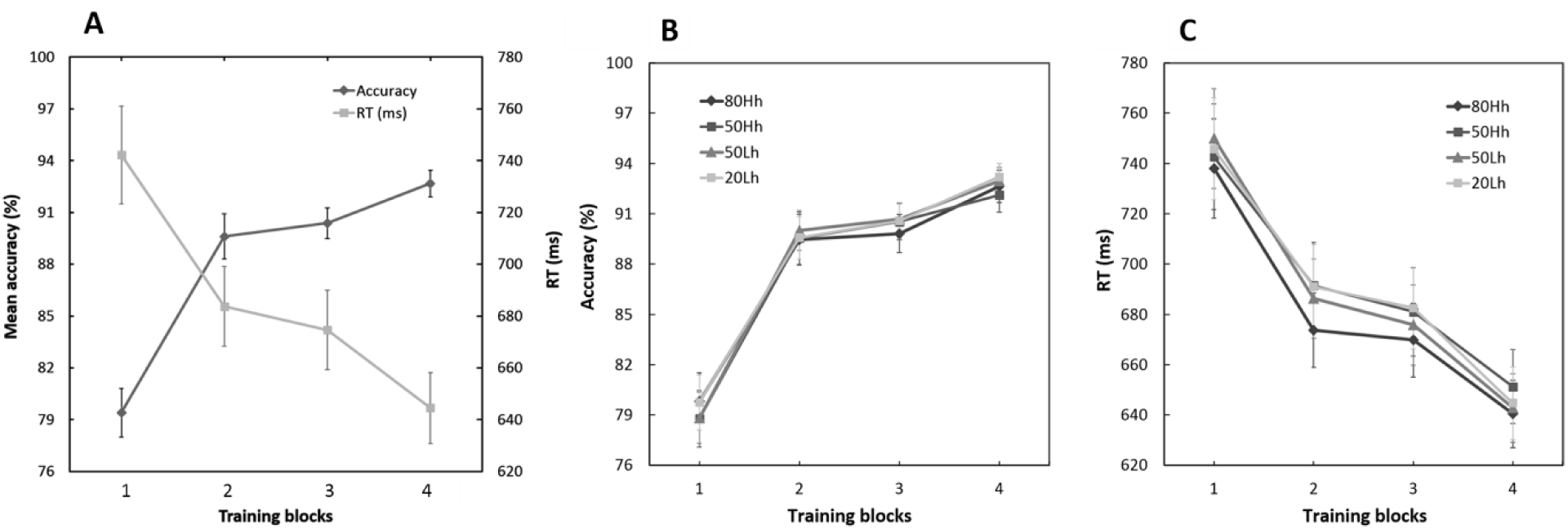
Training performance. (A) Learning profile across the reward-based training sessions: mean accuracy (mean ± SEM) and mean reaction time (RT; mean ± SEM) plotted across four consecutive blocks (two blocks per day). (B) Accuracy (mean ± SEM) and (C) RT (mean ± SEM) as a function of Reward bias (80Hh, 50Hh, 50Lh, 20Lh) across blocks.

To test for learning effects across training, we conducted a two-way repeated-measures ANOVA on ACC with factors block (1-4) and reward bias (80Hh, 50Hh, 50Lh, 20Lh; values indicate the probability of high reward on correct trials, with Hh/Lh denoting the high-/low-reward hemifield, see Methods). As shown in Figure 1A, ACC increased significantly across blocks (*F*(3,117) = 72.221, *p* < 0.001, partial η² = 0.649). However, ACC was not modulated by reward bias (main effect: *F*(3,117) = 0.231, *p* = 0.875) and there was no block × reward bias interaction (*F*(9,351) = 0.554, *p* = 0.834) (Figure 1B). A corresponding repeated-measures ANOVA on RTs showed a significant decrease across blocks (*F*(3,117) = 24.650, *p* < 0.001, partial η² = 0.387), with no evidence for a main effect of reward bias (Figure 1C).

When participants were queried about the reward amount received on each trial, mean accuracy on this reward report was 75.65 ± 2.70 % and 76.31 ± 2.87 % on training sessions 1 and 2, respectively.

#### Baseline and Test sessions

We first evaluated performance during the baseline session (Day 1). In trials with a single target (single target condition), ACC was analyzed using a two-way repeated-measures ANOVA with factors spatial location (1-8) and target type (letter vs. digit). ACC varied significantly across spatial locations (*F*(7, 273) = 36.32, *p* < .001, partial η² = .49), with higher ACC at more lateralized locations relative to locations closer to vertical midline. ACC also differed by target type (*F*(1, 39) = 54.47, *p* < .001, partial η² = .58), with higher ACC for digits than letters (Figure 2A). An interaction between target type and spatial location was not found.

**Figure 2:**
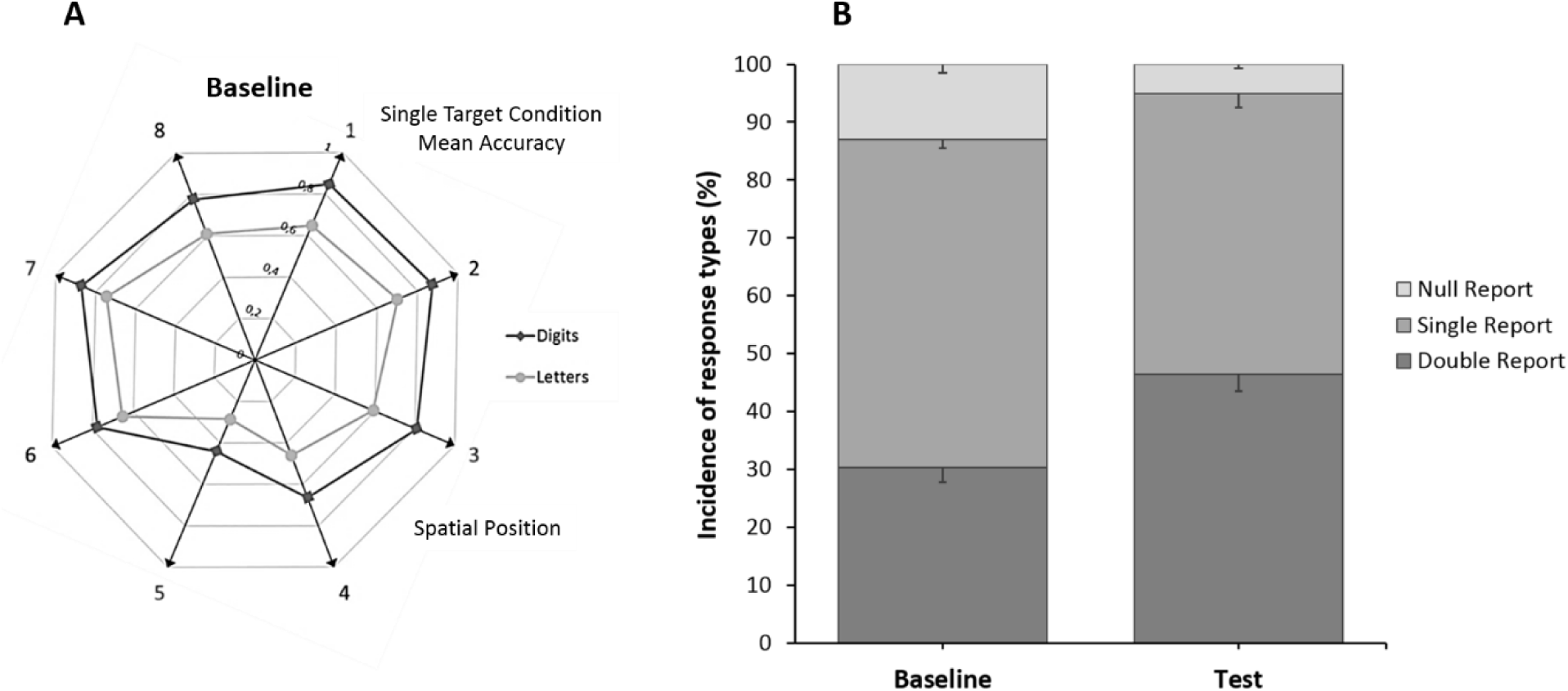
Baseline and test performance of visual search task. (A) Baseline accuracy in the single-target condition. Polar plot showing mean accuracy as a function of spatial location (1–8) and target type (digits vs. letters). (B) Double-target condition response distributions at baseline and test. Mean proportion (mean ± SEM) of double, single, and null reports for the baseline (left bar) and test (right bar) sessions.

#### Double target condition

In the double target condition, participants could correctly report both targets (double report), one target (single report), or neither target (null report). During the baseline session, the mean proportions (± SEM) were 30.26 ± 2.53% for double reports, 56.74 ± 1.50% for single reports, and 13.00 ± 1.45% for null reports. At test (Day 7), *t*-Tests revealed a significant increase in the incidence of double reports relative to baseline (Day 1) (16.11 ± 1.37 %; *t*(39) = 11.761, p < 0.001, r = 0.879), and a significant decrease in the single reports (−8.17 ± 1.51 %; *t*(39) = −5.421, *p* < 0.001, r = 0.791), as well as the null reports (−7.94 ± 1.04 %; *t*(39) = −7.608, *p* < 0.001, r = 0.749) (Figure 2B).

#### Single report during double target condition

Performance in the double target condition was further examined for single report trials as a function of the spatial locations at which the two targets appeared. We selected trials in which one target was presented at a high reward location and the other at a low reward location, yielding four location-pair combinations: 80Hh-20Lh, 80Hh-50Lh, 50Hh-20Lh, and 50Hh-50Lh. By comparing test versus baseline, we tested the prediction that reward based training produces long-lasting alterations in spatial priority maps, conferring a relative advantage to targets presented at highly rewarded compared with poorly rewarded locations (Chelazzi et al., 2014). Therefore, we focused first on the critical 80Hh-20Lh trials. We computed Δ probability as report 80Hh – report 20Lh and tested whether this bias increased from baseline to test. Δ probability did not differ significantly between sessions (Δ probability = −1.895; *t*(39) = −0.800, *p* = 0.429), with the numerical trend opposite to the predicted direction. No significant baseline-to-test change in Δ probability was observed for the other location-pair combinations (Figure 3A).

**Figure 3:**
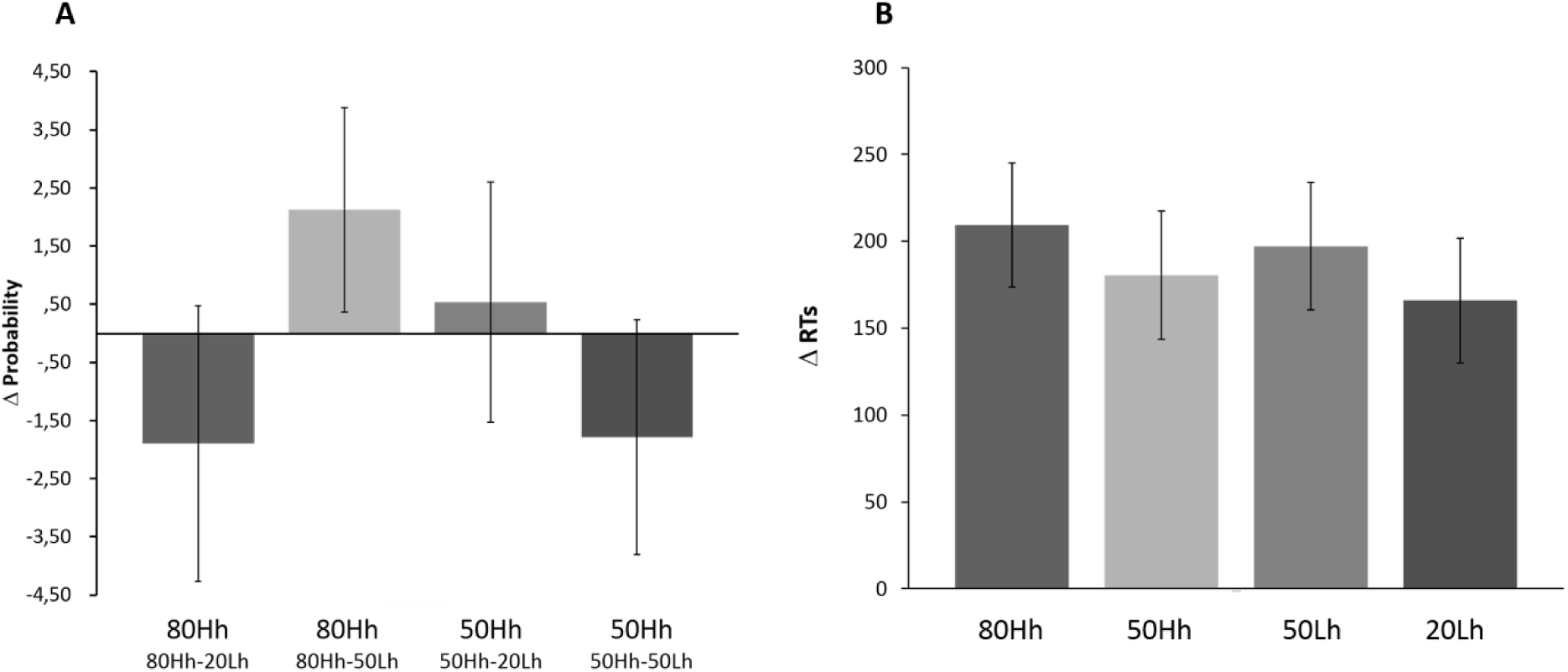
Double-target condition performance by reward-biased location. (A) Change in selection bias (Δ probability) from baseline to test for single-report trials, computed as the difference in the probability of reporting the target presented at the higher-reward location relative to the lower-reward location for each location-pair type. (B) Change in reaction time (RT; test − baseline) on single-report trials as a function of the reward-biased location of the reported target (80Hh, 50Hh, 50Lh, 20Lh).

We additionally analyzed RTs for single-report trials in the double target condition as a function of the reward biased location of the correctly reported target. Because the task was not speeded, RTs outside 200-4000 ms were excluded. A two-way repeated-measures ANOVA with factors session (baseline vs. test) and reward bias (80Hh, 50Hh, 50Lh, 20Lh) showed a significant main effect of session, with faster RTs at test relative to baseline (*F*(1,39) = 35.392, *p* < 0.001, partial η² = .476). There was no main effect reward bias (*F*(3,117) = 0.330, *p* = 0.803) and no session × reward bias interaction (*F*(3,117) = 0.875, *p* = 0.456), indicating that the RT improvement was not selective to the reward manipulation (Figure 3B).

#### Double report during double target condition – what was reported first?

On trials in which both targets were correctly reported (double reports), we examined whether report order showed a bias when the two targets appeared at the most extreme reward-biased locations (80Hh vs. 20Lh). Specifically, we tested whether participants’ first response was more likely to correspond to the target presented at the high-reward (80Hh) versus low-reward (20Lh) location, and whether this tendency differed between baseline and test. We counted how many of their first presses were in the 80Hh position, how many in the 20Lh position in the baseline and how many in the test session. T-tests showed no significant baseline-to-test change in either count, indicating no reliable shift in the report order bias toward the highly rewarded location.

### EEG (ERPs)

#### Training Sessions - After Feedback

The Feedback-Related Negativity (FRN) is a fronto-central, negative-going ERP component typically peaking ∼200–350 ms after feedback onset and commonly larger (more negative) following unfavorable outcomes. We quantified the FRN using three common approaches and tested effects of feedback valence, reward magnitude, and training progression (i.e., data from the two training sessions were binned into four consecutive blocks) over fronto-central/central midline electrodes (Fz, FCz, Cz, CPz; anteriority refers to the electrodes’ front-to-back position along the midline).

#### FRN and feedback valence (correct vs. incorrect)

Using mean FRN amplitude around the peak, a repeated-measures ANOVA with factors valence (correct, incorrect), anteriority (Fz, FCz, Cz, CPz), and block (1–4) showed significant main effects of valence (*F*(1,31) = 7.952, *p* = .008), anteriority (*F*(3,93) = 18.921, *p* < .001), and block (*F*(3,93) = 11.162, *p* < .001), as well as a valence × block interaction (*F*(3,93) = 12.573, *p* < .001). Specifically, FRN amplitudes changed across blocks for correct feedback, whereas this time course was not observed for incorrect feedback (Figure 4). Results were broadly consistent across alternative FRN quantification methods. With a base-to-peak measure relative to the preceding P2, the same ANOVA yielded significant main effects of valence (*F*(1,31) = 24.012, *p* < .001), anteriority (*F*(3,93) = 9.922, *p* < .001), and block (*F*(3,93) = 12.947, *p* < .001), with significant interactions for anteriority × block (*F*(9,279) = 4.265, *p* < .001) and valence × anteriority × block (*F*(9,279) = 2.126, *p* = .028). Using the Yeung–Sanfey scoring approach, significant main effects of valence (*F*(1,31) = 75.481, *p* < .001) and anteriority (*F*(3,93) = 12.063, *p* < .001) were observed, along with a valence × anteriority interaction (*F*(3,93) = 10.514, *p* < .001), whereas the effect of Block was not significant.

**Figure 4:**
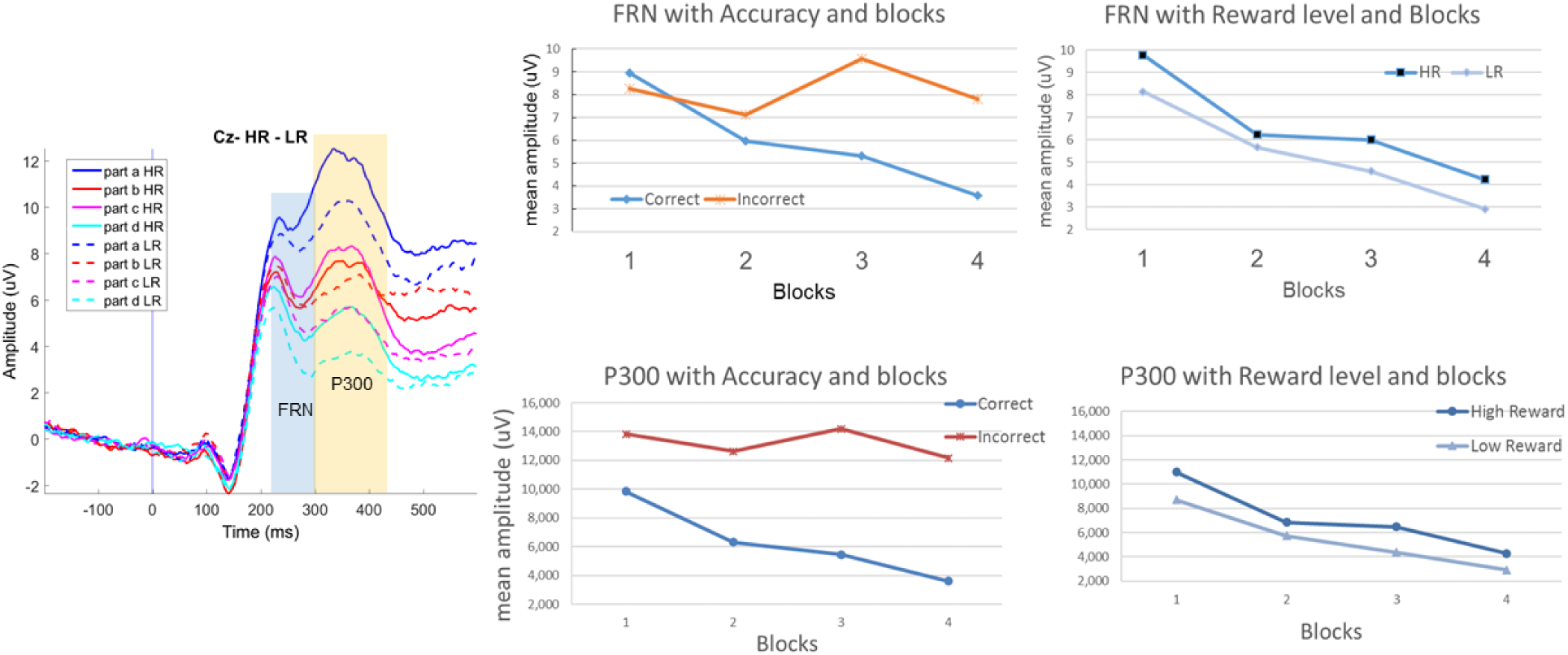
Feedback-locked ERPs during training. Left: grand-average waveforms showing feedback-related FRN and P3 across training blocks. Right: mean FRN (top) and mean P3 (bottom) amplitudes plotted as a function of feedback accuracy (correct vs. incorrect) across blocks, and as a function of reward magnitude (high vs. low) across blocks.

#### FRN and reward magnitude (high vs. low reward; correct trials)

To assess sensitivity to reward magnitude, we analyzed correct-feedback trials only and ran a repeated-measures ANOVA with factors reward level (high, low), anteriority (Fz, FCz, Cz, CPz), and block (1–4). For mean FRN amplitude, there were significant main effects of reward level (*F*(1,31) = 15.868, *p* < .001), anteriority (*F*(3,93) = 20.254, *p* < .001), and block (*F*(3,93) = 59.197, *p* < .001), with FRN amplitudes more negative for low than high reward. Interactions were significant for reward level × anteriority (*F*(3,93) = 3.057, *p* = .032) and anteriority × block (*F*(9,279) = 3.353, *p* = .001). When FRN was quantified using the base-to-peak method, main effects were observed for anteriority (*F*(3,93) = 6.088, *p* = .01) and block (*F*(3,93) = 40.600, *p* < .001), but not for reward level; the only significant interaction was anteriority × block (*F*(9,279) = 2.259, *p* = .019). With the Yeung–Sanfey method, significant interactions were found for reward level × anteriority (*F*(3,93) = 10.801, *p* < .001) and anteriority × block (*F*(9,279) = 2.151, *p* = .026). Follow-up analyses by electrode indicated a reward-level effect at CPz (*F*(1,31) = 7.905, *p* = .008) and a block effect at FCz (*F*(3,93) = 2.841, *p* = .042).

Overall, FRN was sensitive to both feedback valence and reward magnitude and showed systematic modulation across training blocks (see Figure 4).

#### P3 and feedback valence (correct vs. incorrect)

To test whether feedback-locked P3 amplitudes differed for correct versus incorrect outcomes and how this varied over training, we analyzed mean P3 amplitude with a repeated-measures ANOVA with factors accuracy (correct, incorrect), anteriority (Fz, FCz, Cz, CPz), and block (1–4). This ANOVA showed significant main effects of accuracy (*F*(1,31) = 60.432, *p* < .001), anteriority (*F*(3,93) = 12.614, *p* < .001), and block (*F*(3,93) = 12.694, *p* < .001). Significant interactions were also observed for anteriority × accuracy (*F*(3,93) = 12.771, *p* < .001), anteriority × block (*F*(9,279) = 3.344, *p* = .001), and accuracy × block (*F*(3,93) = 6.711, *p* < .001). To account for overlap with the preceding negative deflection (FRN), we also quantified P3 using a base-to-peak measure. This analysis again yielded significant main effects of accuracy (*F*(1,31) = 107.35, *p* < .001) and anteriority (*F*(3,93) = 4.180, *p* = .008), but not block. Significant interactions remained for anteriority × accuracy (*F*(3,93) = 7.052, *p* < .001) and anteriority × block (*F*(9,279) = 3.671, *p* = .001).

#### P3 and reward magnitude (high vs. low reward)

We next tested sensitivity to reward magnitude by analyzing correct-feedback trials as a function of reward level (high, low), anteriority (Fz, FCz, Cz, CPz), and block (1–4). Mean P3 amplitude showed significant main effects of reward level (*F*(1,31) = 48.453, *p* < .001), anteriority (*F*(3,93) = 12.478, *p* < .001), and block (*F*(3,93) = 39.282, *p* < .001). P3 amplitudes were larger for high than low reward (high: 7.161 ± 0.858; low: 5.445 ± 0.790) and decreased across blocks (block 1: 9.855 ± 0.980; block 2: 6.299 ± 0.878; block 3: 5.438 ± 0.928; block 4: 3.618 ± 0.769), with the largest amplitudes at Cz. The reward level × block interaction was significant (*F*(3,93) = 3.927, *p* = .011), indicating a block-related decrease for each reward level (Figure 4). Anteriority × block was also significant (*F*(9,279) = 3.401, *p* = .001), whereas anteriority × reward level was not. Using the base-to-peak measure to reduce FRN overlap, the ANOVA again showed main effects of reward level (*F*(1,31) = 7.007, *p* = .031), anteriority (*F*(3,93) = 7.644, *p* < .001), and block (*F*(3,93) = 4.957, *p* = .003). In this scoring, reward level × block was not significant, whereas reward level × anteriority (*F*(3,93) = 4.730, *p* = .004) and anteriority × block (*F*(9,279) = 2.859, *p* = .003) were significant.

Thus, feedback-locked P3 amplitudes reliably tracked both outcome accuracy and reward magnitude and showed systematic modulation across training blocks, with the spatial distribution of these effects varying across the midline electrodes.

#### Training Sessions - After Stimulus (target)

To assess short-term reward effects on target processing during training, we analyzed target-locked ERPs for the components shown in Figure 5. Unless otherwise noted, repeated-measures ANOVAs included anteriority (Fz, FCz, Cz, CPz, Pz, POz, Oz), reward bias (80Hh vs. 20Lh), and block (1–4).

**Figure 5:**
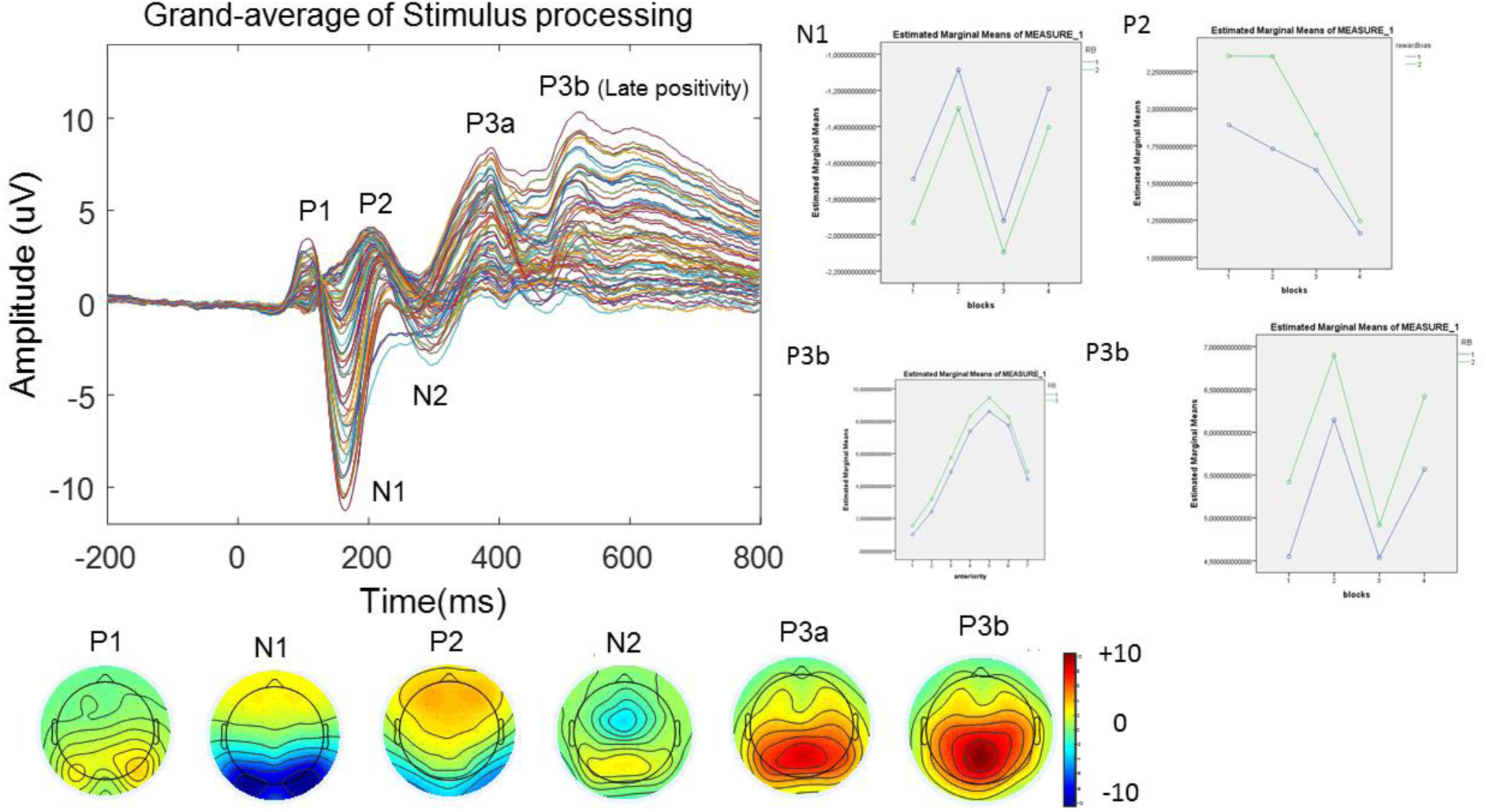
Stimulus-locked ERPs during training. Grand-average target-evoked waveforms and corresponding scalp topographies for each identified component.

##### P1 (P100)

No significant effects were observed for P1.

##### N1 (N100)

The N1 showed significant main effects of anteriority (*F*(6,198) = 68.078, *p* < .001), reward bias (*F*(1,33) = 5.851, *p* = .021), and block (*F*(3,99) = 7.115, *p* < .001), as well as an anteriority × block interaction (*F*(18,594) = 9.751, *p* < .001). Across blocks, N1 amplitudes were consistently larger for targets appearing at the low-reward location (20Lh) relative to the high-reward location (80Hh). Given the scalp distribution of the N1 (Figure 5), we analyzed parieto-occipital sites (PO3, PO4, PO7, PO8). This analysis yielded main effects of anteriority (*F*(3,99) = 4.256, *p* = .007), reward bias (*F*(1,33) = 4.682, *p* = .038), and block (*F*(3,99) = 22.519, *p* < .001), with no significant interactions.

##### N2 (N200)

For the N2, effects were assessed at CPz and Pz. A repeated-measures ANOVA with factors electrode (CPz, Pz), reward bias (80Hh vs. 20Lh), and block (1–4) showed significant main effects of electrode (*F*(1,25) = 19.969, *p* < .001) and reward bias (*F*(1,25) = 4.710, *p* = .040). N2 mean amplitude was more negative for targets at the low-reward location (20Lh) than at the high-reward location (80Hh).

##### Late positivity (P3b)

A later centro-parietal positivity (P3b/late positivity) showed significant main effects of reward bias (*F*(1,26) = 5.729, *p* = .024) and anteriority (*F*(6,156) = 34.375, *p* < .001) and an anteriority × block interaction (*F*(18,468) = 5.019, *p* < .001). Mean amplitude of this late positivity was larger for targets at the low-reward location (20Lh) than at the high-reward location (80Hh).

Thus, during training, reward-biased locations modulated stimulus-locked processing from early sensory-perceptual stages (N1) through later selection/evaluative stages (N2 and late positivity), whereas the earliest P1 component showed no detectable reward-related effects.

#### Baseline and Test sessions

##### Double target condition: ERPs by report outcome (double vs. single vs. null)

ERP analyses focused on midline electrodes, where effects were maximal over centro-parietal sites (for example CPz; Figure 6). Grand-average waveforms in the double-target condition indicated systematic differences in both the N2 and P3 time ranges as a function of report outcome: P3 amplitudes were largest for trials on which participants reported both targets, intermediate for single reports, and smallest for null reports. This ordering was evident in both sessions, alongside an overall reduction in P3 amplitude at test relative to baseline (Figure 6).

**Figure 6:**
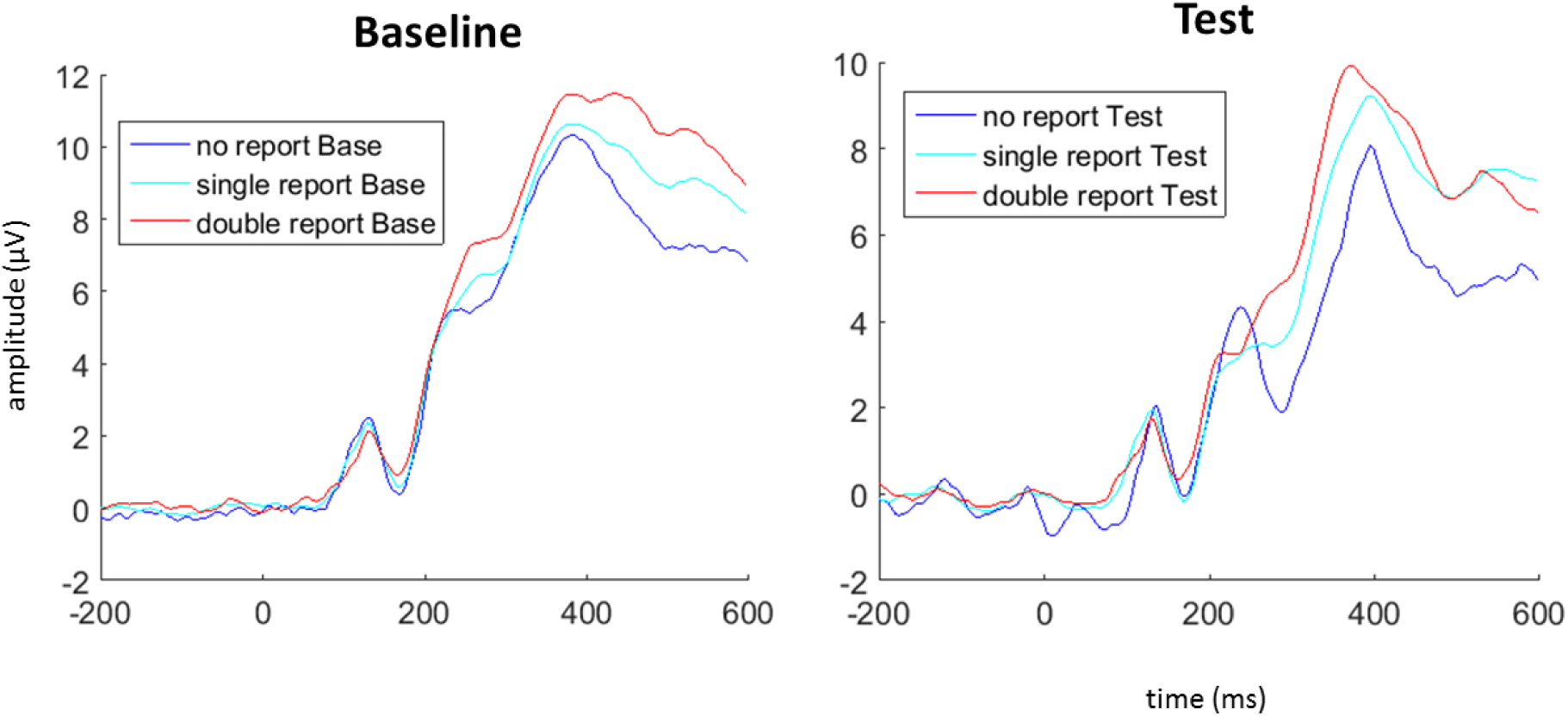
Double-target condition ERPs by report outcome. Grand-average stimulus-locked waveforms at CPz for double, single, and null report trials, shown separately for the baseline (left) and test (right) sessions.

These observations were confirmed statistically. For the N2 mean amplitude, a repeated-measures ANOVA with factors report (double, single, null), session (baseline, test), and anteriority showed significant main effects of anteriority (*F*(6,180) = 3.730, *p* = .003), report (*F*(2,60) = 3.207, *p* = .047), and session (*F*(1,30) = 83.447, *p* < .001), as well as significant interactions for anteriority × session (*F*(6,180) = 22.185, *p* < .001) and report × session (*F*(2,60) = 3.654, *p* = .032). For the P3 mean amplitude, a corresponding ANOVA with factors report, session, and anteriority (AFz, Fz, FCz, Cz, CPz, Pz, POz, Oz) revealed significant main effects of anteriority (*F*(7,210) = 51.900, *p* < .001), report (*F*(2,60) = 12.396, *p* < .001), and session (*F*(1,30) = 45.273, *p* < .001), with significant interactions for anteriority × session (*F*(7,210) = 23.454, *p* < .001) and report × session (*F*(2,60) = 3.822, *p* = .027). Thus, centro-parietal activity in the N2/P3 time ranges scaled with trial outcome (successful report of both targets vs. partial vs. no report) and differed between baseline and test, with P3 amplitudes overall reduced at test.

##### Double-target condition: double-report trials (baseline vs. test)

Figure 7 shows grand-average ERPs for the baseline and test sessions restricted to trials in which participants correctly reported both targets (double reports). In this subset, baseline vs. test differences were evident in both the N2 and P3 time ranges. For the N2, a repeated-measures ANOVA with factors session (baseline, test) and anteriority (Fz, FCz, Cz, CPz, Pz, POz) showed main effects of anteriority (*F*(5,150) = 3.243, *p* = .008) and session (*F*(1,30) = 44.795, *p* < .001), as well as an anteriority × session interaction (*F*(5,150) = 12.984, *p* < .001). Post-hoc paired t-tests indicated significant baseline–test differences at Fz, FCz, Cz, and CPz (all *p* < .001) and at Pz (*p* = .001). For the P3, baseline vs. test differences were assessed using paired two-tailed *t*-tests at midline electrodes. These comparisons showed significant session differences at Fpz, AFz, Fz, Cz, CPz, and Pz. Normality checks (Shapiro–Wilk) did not indicate deviations from normality for the analyzed measures.

**Figure 7:**
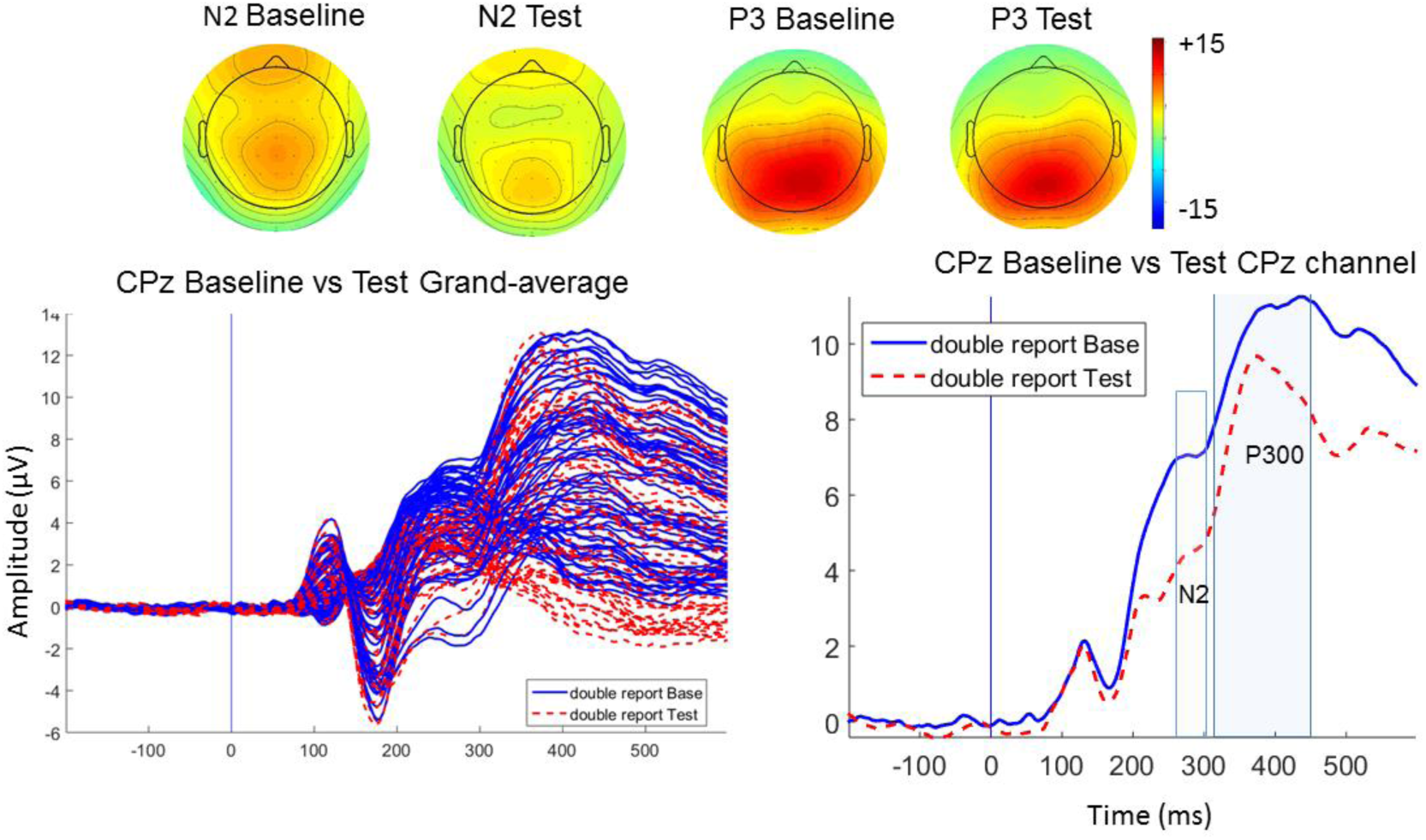
Double target condition, double report trials. Left: 64-channel butterfly plots for the baseline (blue) and test (red) sessions. Right: corresponding grand-average ERP waveforms at CPz.

##### Double-target condition: single-report trials (high vs. low reward locations)

To test for location-specific effects of the reward manipulation at delayed test, we focused on single-report trials in which the reported target appeared at either the highest- (80Hh) or lowest-reward (20Lh) location. Mean N2 amplitude was analyzed with a repeated-measures ANOVA with factors anteriority (AFz, Fz, FCz, Cz), session (baseline, test), and reward bias (80Hh vs. 20Lh). This analysis showed significant main effects of anteriority (*F*(3,87) = 5.881, *p* = .001) and session (*F*(1,29) = 64.042, *p* < .001), but no main effect of reward bias. Importantly, a session × reward bias interaction was observed (*F*(1,29) = 5.987, *p* = .027). Follow-up paired t-tests (Table 1) showed that 20Lh and 80Hh trials differed at test at FCz (*p* = .023) and Fz (*p* = .029), whereas the corresponding comparisons at baseline were not significant (FCz: *p* = .273; Fz: *p* = .366).

**Table 1:** Post-hoc paired-samples t-tests comparing N2 mean amplitude for targets reported at the 80Hh versus 20Lh locations in the double-target condition (single-report trials), shown separately for baseline and test sessions.

| Post-hoc paired sample test | mean | SD | *p* value |
| --- | --- | --- | --- |
| FCz Baseline 20Hh | 4.930 | 5.508 | 0,273 |
| FCz Baseline 80Hh | 4.551 | 5.402 |  |
| FCz Test 20Hh | 0.665 | 5.451 | 0,023 (*) |
| FCz Test 80Hh | 1.581 | 5.408 |  |
| Fz Baseline 20Hh | 4.538 | 5.325 | 0,366 |
| Fz Baseline 80Hh | 4.211 | 5.300 |  |
| Fz Test 20Hh | 0.373 | 5.336 | 0,029 (*) |
| Fz Test 80Hh | 1.518 | 5.513 |  |

Mean P3 amplitude was analyzed with a repeated-measures ANOVA with factors anteriority (AFz, Fz, FCz, Cz, CPz, Pz, POz, Oz), session (baseline, test), and reward bias (80Hh vs. 20Lh). The ANOVA revealed main effects of anteriority (*F*(7,203) = 57.24, *p* < .001), session (*F*(1,29) = 17.30, *p* < .001), and reward bias (*F*(1,29) = 7.543, *p* = .010), as well as an anteriority × session interaction (*F*(7,203) = 21.477, *p* < .001). The session × reward bias interaction was not significant, indicating that the overall reward-bias effect in the P3 time range did not differ reliably between baseline and test.

When restricting analyses to trials in which the two targets appeared specifically at the 80Hh–20Lh location pair, the number of retained trials per condition was low. Median trial counts (± SD) were 13 ± 8 (20Lh) and 15 ± 7 (80Hh) at baseline, and 12 ± 8 (20Lh) and 12 ± 6 (80Hh) at test. Given these low counts (after artifact rejection), we did not conduct reliable statistical comparisons for pair-specific ERPs (see Supplementary Information). Figure 8 shows the ERP waveforms at FCz for these trial pairs.

**Figure 8:**
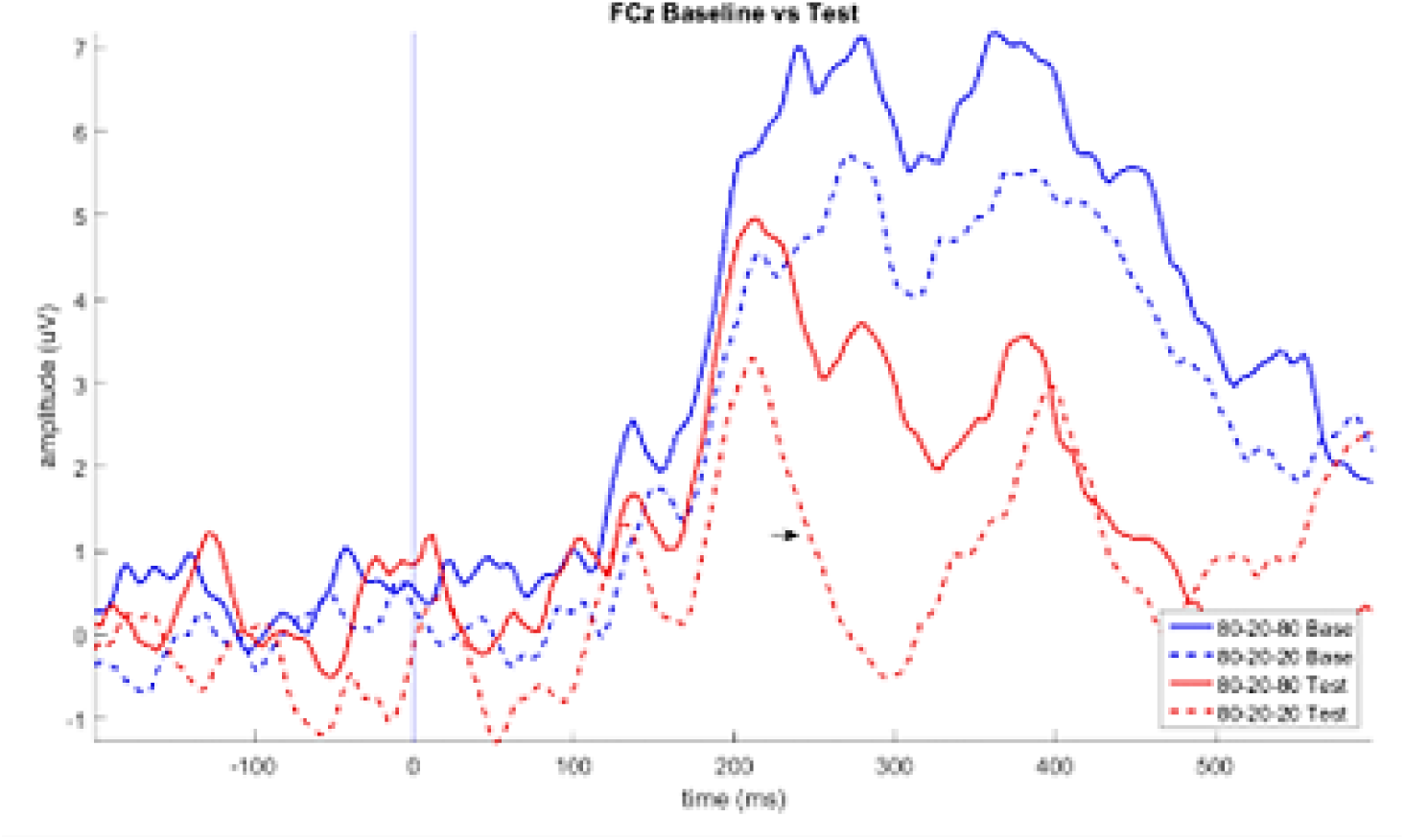
Double-target condition, single-report trials (80Hh–20Lh pair). Grand-average ERP waveforms shown separately by session (baseline vs. test) and reward-biased location (80Hh vs. 20Lh).

### Pupillometry

#### Pupil responses during training - after feedback

To assess sensitivity to reward magnitude, we analyzed feedback-locked pupil responses using the same baseline-corrected 750 ms window (1000–1750 ms post-feedback onset). A repeated-measures ANOVA with factors block (1–4) and reward level (high, low) showed main effects of block (*F*(3,108) = 5.600, *p* = .001) and reward level (*F*(1,36) = 21.076, *p* < .001), indicating that pupil dilation decreased over blocks and was larger following high- versus low-reward feedback (Figure 9B). The reward level × block interaction was also significant (*F*(3,108) = 4.145, *p* = .008).

**Figure 9:**
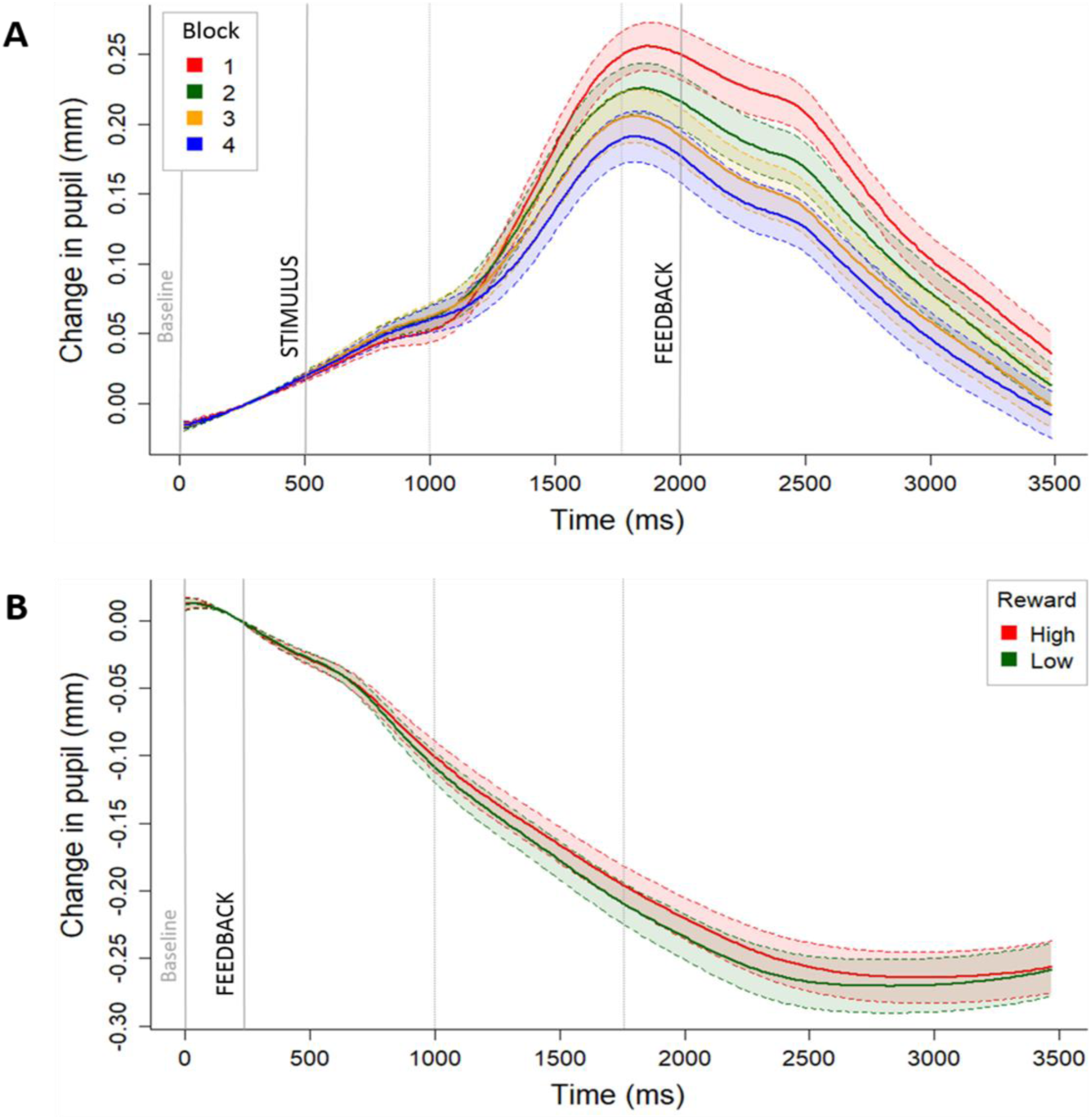
Task-evoked pupil responses during training. Baseline-corrected pupil change (mm) following (A) stimulus (target) onset, shown separately for the four training blocks, and (B) feedback onset, shown separately for low- and high-reward feedback.

#### Pupil responses during training - after stimulus (target)

To test whether pupil diameter reflected short-term reward-related differences across spatially biased locations, we analyzed stimulus-locked pupil responses during training. Baseline-corrected mean pupil change was computed over a 750 ms window (1000–1750 ms post-stimulus onset). A repeated-measures ANOVA with factors block (1–4) and reward bias (80Hh, 50Hh, 50Lh, 20Lh) showed a significant main effect of block (*F*(3,108) = 28.533, *p* < .001), with pupil responses decreasing across training blocks (Figure 9A). There was no main effect of reward bias and no block × reward bias interaction. Thus, pupil responses showed a general decrease across training blocks and reliably differentiated feedback reward magnitude, while showing no evidence of location-specific modulation during target processing.

## Discussion

This study applied EEG and pupillometry to a multi-session spatial reward-learning paradigm previously used to argue that biased reward schedules can induce long-lasting changes in spatial priority that generalize to a different task context. Our results provide evidence for a dissociation: during the rewarded training task, reward evaluation signals were robust and multimodal (feedback-locked ERPs and pupil dilation), and target processing showed location-related ERP modulations. In contrast, evidence that these learned spatial values transfer to a separate task after a multi-day delay was less consistent, with only a limited N2 modulation detectable under conditions of low trial counts for the critical high-vs-low reward pairings. This pattern sharpens the interpretation of spatial value-driven attention by separating intact *reinforcement learning* from the *persistence* and *generalization* of spatial attentional biases.

This constrains claims that spatial reward associations automatically consolidate into persistent, context-general priority-map changes (Chelazzi et al., 2014; Della Libera et al., 2017). More recent work and synthesis in the broader value-driven attention literature suggest that the robustness and generalizability of learned value biases can depend on factors such as uncertainty, individual differences, and task context, often yielding effects that are reliable in some settings but fragile or heterogeneous in others (Anderson & Britton, 2019; Anderson et al., 2021; Garre-Frutos et al., 2024; Stanković, 2025; Theeuwes, 2019). In that light, our behavioral pattern is consistent with the possibility that spatial value-driven biases are more sensitive to contextual change (task demands, stimulus set, response regime) than the well-established feature-based value effects (Anderson, 2013; Anderson et al., 2011; Hickey et al., 2010; Libera & Chelazzi, 2006).

### Robust reward learning during training despite weak long-term behavioral transfer

Across training, feedback-locked ERPs showed strong sensitivity to outcome properties and time-on-task. FRN and feedback-locked P3 reliably differentiated feedback valence (correct vs. incorrect) and showed systematic modulation across blocks, consistent with classic interpretations of these components as indexing early outcome evaluation and later updating/attentional allocation to motivationally significant feedback (Holroyd & Coles, 2002; San Martín, 2012). The observed modulation of these signals across blocks is compatible with changing learning demands and/or changing utility of feedback as performance stabilizes. P3 sensitivity to reward magnitude and/or valence is a common finding in reward-learning and task-monitoring ERP work, and is often interpreted as reflecting updating of internal context, salience, and the allocation of processing resources to feedback that carries learning-relevant information. This dissociation matters theoretically. Within the “selection history” framework, reward learning is one source of history-dependent bias that can compete with goals and physical salience (Anderson et al., 2021; Theeuwes, 2019). Our results suggest that the existence of strong neural valuation signals during learning is not sufficient to guarantee a stable, task-general spatial bias that persists after reinforcement is removed.

Pupillometry converged with these interpretations: feedback-evoked pupil dilation differentiated reward magnitude and tended to decrease with time-on-task (Preuschoff et al., 2011; Van Slooten et al., 2018). Such a pattern is consistent with reduced effort/arousal demands as task structure becomes more predictable, alongside preserved sensitivity to motivational outcomes (Mathôt, 2018; Van der Wel & Van Steenbergen, 2018). Together, the multimodal outcome-evaluation results strengthen a key inference: participants encoded and responded to the reward structure during training, so the later absence of strong transfer cannot be attributed to a trivial failure to learn the reward contingencies (Holroyd & Coles, 2002; San Martín, 2012).

### Reward history modulates target processing during training

Stimulus-locked ERPs during training showed location-dependent modulations spanning later processing stages (notably N2-range activity and a late positivity). This supports the idea that spatial reward learning influences not only feedback evaluation but also subsequent target processing during active learning, consistent with accounts in which learned value biases attentional competition by modulating priority signals and downstream evaluative/control processes (Itthipuripat et al., 2019; Krauzlis et al., 2014).

A notable feature of the training ERPs is that targets at the low-reward location (20Lh) evoked larger responses than targets at the high-reward location (80Hh), spanning an attentional/sensory N1 modulation and later N2-range and late positive activity. One interpretation is compensatory selection: if reward learning biases spatial priority toward high-value locations, selecting a target at a low-value location may require greater attentional amplification and post-perceptual selection effort, yielding larger N1/N2 and a larger late positivity. The N1 is an early occipito-temporal component that is reliably enhanced when attention amplifies processing in higher-tier visual cortex, making it a sensitive marker of attentional gain during perceptual encoding (Hillyard & Anllo-Vento, 1998; Luck et al., 2000). N2-range activity is often modulated by stimulus discrimination difficulty, competition/selection demands, and cognitive control processes (Folstein & Van Petten, 2008).

However, several alternative interpretations warrant consideration. One possibility is surprise/expectancy violation: low-reward locations may be less expected, such that selecting a 20Lh target elicits stronger evaluative responses because it deviates from learned value-based expectations, consistent with predictive coding accounts in which unexpected events evoke enhanced neural responses (Friston, 2005; Rao & Ballard, 1999). A second possibility is conflict/control recruitment: selecting a target at a low-value location when a high-value alternative is concurrently available may increase competition between learned value bias and current selection, engaging control mechanisms often associated with N2-family activity and conflict monitoring (Botvinick et al., 2001; Yeung et al., 2004).

The present design does not distinguish these mechanisms definitively, but the consistent low > high pattern across components indicates that reward history altered target processing during learning, even though delayed behavioral transfer was weak. Notably, this pattern contrasts with typical N2pc results showing larger amplitudes for high-reward distractors (Itthipuripat et al., 2015; Qi et al., 2013), suggesting the effect may be specific to target (rather than distractor) processing or the spatial nature of the manipulation. Future studies using designs that independently manipulate target and distractor value, and that include measures of surprise and conflict, could help dissociate these accounts.

### Behavioral results at delayed test: context dependence and measurement constraints

Using the behavioral logic of the original report (Chelazzi et al., 2014), we did not find reliable evidence that highly rewarded locations retained a durable advantage in the delayed test task. Consistent with this constraint, a conservative interpretation is that spatial value learning is more context-dependent and/or more heterogeneous across individuals than strong versions of the “persistent spatial priority” hypothesis imply. Value-driven attentional biases frequently show context gating, strong within a learned context but attenuated when task demands, stimulus sets, or spatial context changes (Anderson & Britton, 2019; Anderson & Kim, 2018). In particular, spatial value learning may depend on the representational format of space: biases can be robust when linked to meaningful spatial layouts or scenes, yet weaker when “space” is abstracted to repeated positions without stable contextual landmarks. On this view, the multi-day reward schedule can produce strong learning signals, but the learned task-specific selection policy may remain tied to the training task’s specific perceptual-motor demands and feedback structure rather than automatically “rewiring” a general priority map for those locations.

Independent of theory, the delayed-test behavioral metric in this paradigm is statistically fragile: the most diagnostic comparisons rely on relatively rare trial types (e.g., single-report trials in high-vs-low target pairings), and improved performance at test can further reduce those counts and consequently number of trials. This fragility is compounded by recent evidence that value-driven attentional capture measures can show variable reliability depending on analytic choices and often require more trials than are typically collected. Garre-Frutos et al. (2024) demonstrated that reliability varies substantially across analytic specifications, with some pipelines yielding reliability below thresholds needed for individual-differences work. Similarly, Stanković (2025) reported substantial heterogeneity in value-driven capture effect sizes across 52 experiments and identified multiple methodological moderators. These findings suggest that null effects in value-driven attention paradigms should be interpreted cautiously, as they may reflect limited power and measurement reliability rather than a true absence of reward-history influences.

### Delayed neural effects: an N2 trace of reward history with limited reliability

In the baseline/test task, the clearest reward-history effect was an N2 modulation over frontal sites in the test session but not in baseline. Frontal N2 activity is often linked to control-related operations such as conflict monitoring, response competition, and the need for increased top-down selection when alternatives compete (Botvinick et al., 2001; Folstein & Van Petten, 2008). In this context, the pattern is compatible with the idea that learned spatial value can persist as a biasing signal that changes the control demands of selection at test. For example, by increasing monitoring when selecting against a previously higher-value competitor, or by engaging gating mechanisms that regulate which location wins competition under uncertainty (Anderson et al., 2021; Theeuwes, 2019). The finding that N2 (rather than P3 or earlier components) shows this delayed effect is theoretically interesting. If reward history produced a persistent change in spatial priority, one might expect modulation of components associated with attentional allocation (e.g., P3) or early sensory processing (e.g., N1). The absence of such effects, combined with the presence of N2 modulation, suggests that the persistent trace of reward learning may be expressed primarily in control/evaluation dynamics rather than in priority per se. This interpretation aligns with the conflict monitoring account of N2 function and suggests that reward history may influence how selection is controlled rather than simply what is selected. Yet, this interpretation remains tentative because the critical high-versus-low comparisons were supported by relatively low per-subject trial counts after cleaning, which reduces the reliability of ERP estimates and increases sensitivity to scoring choices and window definitions (Luck, 2014). The delayed N2 effect is best viewed as suggestive evidence for a persistent reward-history trace in control/selection-related processing, but one that requires higher-powered designs (e.g., more trials) and reliability-aware, pre-specified scoring to support stronger claims.

### Implications for theories of value-driven spatial attention

The present findings refine the link between reinforcement learning and attentional priority. Training-phase data strongly support intact outcome evaluation and learning (FRN/P3 and pupil; (Cole et al., 2022; Holroyd & Coles, 2002; Megemont et al., 2022; San Martín, 2012)), and show that reward history modulates target-evoked processing during the rewarded task, consistent with value signals biasing spatial priority computations within attention networks (Chelazzi et al., 2014; Itthipuripat et al., 2019; Krauzlis et al., 2014). Yet these learning-related signals were not accompanied by a robust, task-general behavioral bias after a multi-day delay, consistent with the idea that value-driven biases can be context-sensitive (Anderson, 2015). One mechanistic interpretation is policy gating: learned value influences selection when the test context reinstates key control/motivation contingencies from training, but expression is attenuated when stimuli or motivation change. Another possibility is that spatial value learning persists but is expressed more readily in control/evaluation dynamics (e.g., N2-range activity) than in overt performance, especially when behavioral measures are compressed by high overall accuracy or by reduced speed pressure (Folstein & Van Petten, 2008; Heitz, 2014; Ratcliff & McKoon, 2008). Distinguishing these possibilities will require designs that jointly optimize behavioral sensitivity and neural reliability at test, with reliability-aware analysis choices (Garre-Frutos et al., 2024; Luck & Gaspelin, 2017; Stanković, 2025).

The potential difference between spatial and feature-based reward learning deserves also consideration. Spatial and feature attention differ in their normalization properties and scope of influence (Ni & Maunsell, 2019). Spatial attention effects are retinotopically specific and scale with stimulus competition, whereas feature attention effects spread globally and do not depend on stimulus number. These differences may translate to reward learning: feature-based reward associations may be encoded more abstractly and thus generalize more readily across contexts, whereas spatial reward learning may remain more tied to specific retinotopic configurations and competitive dynamics. This hypothesis could be tested in future studies that directly compare spatial and feature-based reward learning within the same participants and paradigm.

### Novelty and relation to prior work

To our knowledge, this is the first study to add both EEG and pupillometry to the multi-day, spatially biased reward-learning paradigm, enabling a joint characterization of outcome evaluation during training and reward-history traces after a delayed, context-changed test. Most neurophysiological work on value-driven attention has largely focused on feature-based learning (e.g., rewarded colors) and has typically used single-session designs and selection-related ERP markers (e.g., N2pc) or imaging approaches (Anderson, 2013; Anderson & Britton, 2019; Anderson et al., 2011; Hickey et al., 2010). In this context, multi-component characterization that spans (a) reinforcement-learning/feedback evaluation (FRN/P3), (b) autonomic arousal/effort dynamics (pupil), and (c) stimulus-locked selection/evaluation signals, and then probes for delayed expression under context change, remains uncommon (Anderson & Britton, 2019; San Martín, 2012). Recent work has emphasized that behavioral estimates of value-driven attentional biases can show variable reliability and can depend on analytic choices, underscoring the value of multimodal measures that can dissociate intact reward learning from weak or heterogeneous behavioral transfer (Freichel et al., 2023; Garre-Frutos et al., 2024; Stanković, 2025).

### Limitations and implications for future work

Several factors likely limited sensitivity to delayed behavioral transfer and neural responses: the un-speeded response regime in the baseline/test task may compress RT-based indices and shift emphasis toward accuracy (Heitz, 2014; Ratcliff & McKoon, 2008); the critical 80Hh–20Lh competitive trials yielded low per-subject counts after cleaning, limiting ERP signal-to-noise and the stability of component estimates (Boudewyn et al., 2018; Luck, 2014); task/context change between training and test may engage different attentional policies (Anderson, 2015); and individual differences in reward sensitivity and/or strategy can be substantial in reward-attention paradigms (Della Libera et al., 2017; Stanković, 2025). Future studies could increase diagnostic trial counts and/or redesign the test to preserve competitive trial types (Boudewyn et al., 2018), incorporate computational learning indices that estimate trial-by-trial value updates (e.g., Wilson and Collins (2019)), assess explicit awareness of contingencies given evidence that awareness can shape reward-attention effects in some paradigms, and use reliability-oriented analysis pipelines (e.g., hierarchical models, single-trial EEG metrics) to reduce flexibility and improve reproducibility (Luck & Gaspelin, 2017).

### Translational and cross-domain relevance

Even in the absence of robust long-term behavioral transfer, the training-phase multimodal signals provide objective markers of reinforcement-learning engagement and outcome evaluation, with autonomic responses that track learning-relevant variables such as surprise/value updating at appropriate timescales (Koenig et al., 2018; Preuschoff et al., 2011; Van Slooten et al., 2018). Robust learning signals during training do not, by themselves, guarantee consolidation or generalization of learned biases to new task contexts. This matters for researchers and practitioners designing reward-based training, neuroadaptive learning systems, and rehabilitation protocols, where success is often evaluated by transfer rather than by learning signals per se. In such settings, neural and autonomic markers can help verify that reinforcement learning is engaged online (Sitaram et al., 2017), while highlighting the need to design training regimes that explicitly target generalization (e.g., by varying context, stimulus sets, and task demands and by incorporating transfer tests as part of the training pipeline) rather than assuming that generalization will automatically follow from strong reward responses (Anderson, 2015).

## Conclusion

Using a multi-day, eight-location spatial reward-learning paradigm with EEG and pupillometry, we provide the first multimodal neurophysiological characterization of the Chelazzi et al. (2014) approach. Training effects were strong: reward contingencies robustly engaged outcome-evaluation signals (FRN/P3 and pupil dilation) and modulated target-evoked processing, indicating effective reinforcement learning and value-dependent influences on attentional processing within the rewarded context. Transfer was weak: applying the original transfer logic, we found no reliable evidence for a persistent spatial attentional bias at delayed test, suggesting limits on how readily spatial value learning consolidates into context-general priority-map changes and consistent with potential heterogeneity in spatial value effects (e.g., Della Libera et al., 2017). Overall, the pattern (i.e., strong reward signals but weak transfer) constrains claims of durable spatial priority-map plasticity across task contexts and motivates future work with higher-powered, reliability-aware designs to determine whether the reduced delayed expression reflects fundamental constraints on spatial priority learning or methodological sensitivity.

## Materials and Methods

### Participants

Forty-three healthy participants were recruited. Three were excluded prior to analysis, leaving a final sample of 40 (mean age = 25.1 years, SD = 3.78; range = 18–36; 22 female). All reported normal or corrected-to-normal vision and no prior experience with similar tasks. Participants provided written informed consent. Compensation was 140 NOK per day for the baseline and test sessions, and 300–500 NOK for training sessions depending on overall accuracy (ACC). The study was approved by the Department of Psychology’s Research Ethics Committee at the University of Oslo.

### Stimuli and Procedure

The stimuli and procedures follow the procedures as in Chelazzi et al. (2014), with minor changes to stimulus color to optimize pupillometric measurement of dilation and constriction. Participants completed four sessions across four separate days: a Baseline session (Day 1), two Training sessions (Days 2–3), and a Test session conducted 4 days after the final Training session. Baseline and Test were identical. Testing took place individually in a sound-attenuated, light-controlled room with constant illumination. Before each task, participants received task-specific instructions and completed 25 practice trials. Baseline and Test lasted ∼1 h each; each Training session lasted ∼1.5 h. Stimuli were presented on a 24-inch BenQ XL2420T LCD monitor (1920 × 1080 pixels; 100 Hz). Viewing distance was 57 cm and head position was stabilized with a chinrest. Tasks were implemented in E-Prime 2.0 Professional (Psychology Software Tools). Responses were collected via a standard keyboard in Baseline/Test and via a response box in Training.

The stimuli used for the Baseline and Test tasks were blue (RGB values: 39, 100, 255) characters (1.2° × 1.2°) displayed on a gray background. Targets comprised eight alphanumeric characters: four capital letters (F, G, M, D) and four digits (2, 4, 7, 9). Distractors were seven non-alphanumeric characters 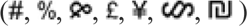, randomized across spatial locations. On each trial, participants searched either for one target (single-target condition; 1 target + 7 distractors) or for two targets (double-target condition; 2 targets + 6 distractors). Trials started with a central blue fixation cross (0.3° × 0.3°; 500 ms), followed by an 8-item stimulus array presented for 70 ms at iso-eccentric positions on an imaginary circle (eccentricity 5°) (Figure 10A). The array was arranged so that each hemifield contained four stimuli, two per quadrant. The eight stimuli were replaced by eight identical masking patterns consisting of overlapping distractors, which remained visible until the participant delivered the behavioral responses. Participants reported up to two detected targets per trial by pressing the corresponding keys for the identified targets, or the spacebar to indicate no target detected. No feedback was provided. The next trial started after a 2500 ms fixation display. The task comprised 640 trials: 192 single-target trials and 448 double-target trials. In the single-target trials, letters and digits occurred with equal probability and each target appeared equally often at each of the eight locations. In the double-target trials, target pairings included letter–letter, digit–digit, and letter–digit combinations; all possible target pairs occurred equally often across all location combinations, balanced to equate spatial relations between the two targets. Conditions were intermixed in random order, and targets were presented equally often at each location.

**Figure 10:**
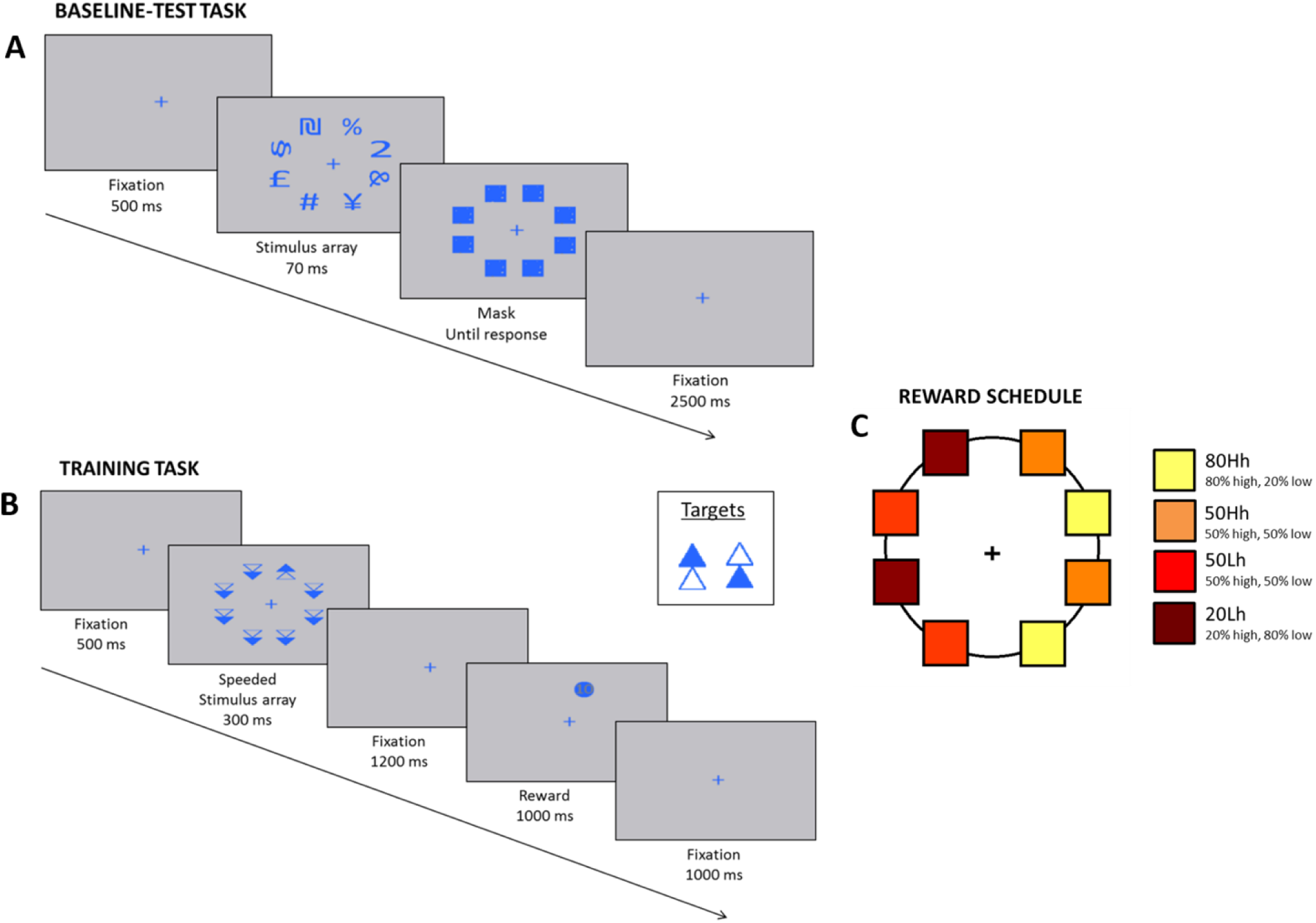
Task designs and reward contingencies. (A) Baseline/Test task trial sequence. (B) Training task trial sequence. (C) Example participant-specific reward schedule showing location-dependent probabilities of high vs. low reward.

In the Training sessions trials began with a central blue fixation cross (0.3° × 0.3°; 500 ms), followed by an 8-item array of simple geometric stimuli (two stacked triangular outlines; 1.2° × 0.7°) presented at the same eight locations as in the Baseline/Test. Each array contained one target (present on every trial) among seven distractors and was displayed for 300 ms, followed by a 1200 ms fixation period to allow pupil diameter to return toward baseline (Figure 10B). Stimuli contained one blue-filled triangle (RGB: 39, 100, 255) and one background-colored triangle (outlined in blue). Distractors were downward-pointing stacked triangles; targets were upward-pointing stacked triangles with the two internal color configurations reversed across target types. Participants were asked to discriminate the target’s internal configuration by reporting the color of the upper triangle as quickly and accurately as possible within 1500 ms of array onset, using two response-box buttons (right index finger for “upper triangle blue”; left index finger for “upper triangle background-colored”). After the response was given, feedback was presented for 1000 ms at the target location. Correct responses received either high (10 points) or low (1 point) feedback; incorrect or missed responses received a red fixation cross. The inter-trial interval was 1000 ms. Each Training session comprised 800 trials (1600 total). The two target configurations occurred equally often and were randomized across trials and locations.

On a subset of participants, reward comprehension was assessed on ∼10% of correct trials by presenting a prompt requiring participants to indicate whether they had received high or low reward using the response box (one button per option). Participants were informed that training compensation depended on performance.

#### Reward contingencies

Reward outcomes were determined by a pre-specified schedule in which reward probability depended on target location, creating a hemifield-level imbalance (Figure 10C). Across participants, one hemifield was designated as the high-reward hemifield (counterbalanced left/right). Within this hemifield, two locations delivered high reward on 80% of correct trials (80Hh) and two locations delivered high reward on 50% of correct trials (50Hh). In the low-reward hemifield, two locations delivered high reward on 20% of correct trials (20Lh) and two locations delivered high reward on 50% of correct trials (50Lh). Thirty-two distinct spatial configurations of these contingencies were generated, and one configuration was assigned to each participant.

### EEG and Pupil recordings

EEG was recorded in all sessions (Baseline, Training, and Test) using a BioSemi ActiveTwo system (BioSemi, Amsterdam, The Netherlands) with 64 active electrodes positioned according to the extended 10–20 system (Oostenveld & Praamstra, 2001) and mounted in an elastic cap (BioSemi). Signals were sampled at 2048 Hz with 24-bit resolution. The EEG was referenced offline to the linked left and right earlobes. Horizontal and vertical electrooculogram (EOG) channels were recorded to monitor eye movements and blinks. The signals were recorded with a sampling frequency of 2048 Hz, and 24 bits resolution. Pupil diameter was recorded during the two Training sessions using an iView X eye-tracking system (SensoMotoric Instruments, SMI) at 60 Hz. To minimize circadian variability in EEG activity, each participant’s sessions were scheduled at approximately the same time of day. Participants were instructed to minimize body and facial movements and to blink naturally while avoiding excessive blinking. A chinrest was used to stabilize head position for pupillometry, and participants were instructed to maintain fixation on the center of the screen. Behavioral accuracy (ACC) and reaction times (RTs) were recorded via E-Prime 2.0 Professional.

### EEG Analysis

EEG preprocessing was performed in MATLAB using EEGLAB (Delorme & Makeig, 2004) and ERPLAB (Lopez-Calderon & Luck, 2014). Continuous data were re-referenced to the average of the left and right earlobe electrodes. For event-related potentials (ERPs) analyses, data were band-pass filtered with a non-causal Butterworth IIR filter (0.1 – 30 Hz; 12 dB/octave, half-amplitude cutoffs) and down sampled to 256 Hz in ERPLAB. Epochs were extracted with a −200 ms pre-event interval, time-locked to stimulus onset (Baseline/Test) or feedback onset (Training). To maximize sensitivity to artifacts, minimally filtered data (DC offset removal only) were visually inspected to identify eye movements, blinks, and muscle activity; contaminated trials were excluded from further analyses. Datasets in which >20% of trials were marked as noisy were excluded, following standard ERP quality-control recommendations (Luck, 2014).

#### Training ERPs time-locked to stimulus onset

Because blink activity was common ∼200–300 ms and after ∼550 ms post-stimulus in Training, an additional ICA decomposition was applied to the stimulus-locked Training datasets to isolate ocular artifacts. Typically, one to two ICA components capturing eye blinks and/or eye movements were removed, and visually rejected trials were retained as excluded (Cohen, 2014). ERP components (P1, N1, P2, N2, P3a, P3b) were identified in each participant’s averaged waveforms based on their temporal order and scalp topography across the 64-channel montage (butterfly plots and topographic maps). Given known inter-individual variability in visually evoked ERP morphology (Woodman, 2010), component latencies were quantified at the single-subject level by measuring each participant’s peak latency for each component for subsequent analyses.

#### Training ERPs time-locked to feedback onset

For feedback-locked ERPs, we quantified the feedback-related negativity (FRN) and the P300. Analyses focused on midline electrodes (Fz, FCz, Cz, CPz), where these components showed maximal activity in the scalp topographies and which are commonly used in the FRN/P300 literature (Cohen et al., 2007; Gehring & Willoughby, 2002; Nieuwenhuis et al., 2004; Yeung & Sanfey, 2004). To assess within-session changes, each training day was split into two blocks of 400 trials. FRN amplitude was quantified in a 250–300 ms window, defined from the grand-average waveform and consistent with prior work. We used three complementary metrics. (1) Mean amplitude in the 250–300 ms window, which is relatively robust to high-frequency noise and modest differences in window placement (Luck, 2014). (2) Peak FRN relative to the preceding positivity (P2), computed as the most negative value in the FRN window referenced to the immediately preceding P2 peak (Hajcak et al., 2006; Janssen et al., 2016; Warren & Holroyd, 2012). (3) Peak-to-peak FRN, computed by referencing the FRN minimum to the average of the surrounding positive peaks (preceding P2 and following P3), to reduce the likelihood that apparent FRN changes reflect shifts in adjacent components (Cohen et al., 2007; Cohen & Ranganath, 2007; Lange et al., 2012; Yeung & Sanfey, 2004). P300 was quantified in a 300–420 ms window using two metrics. (1) Mean amplitude in the 300–420 ms window. (2) Base-to-peak amplitude, computed as the maximum value in 300–420 ms minus the minimum value in the preceding 250–300 ms interval (i.e., the FRN trough), to verify that P300 effects were not driven by baseline shifts or modulation of the preceding negative deflection.

#### Baseline and Test ERPs

For the Baseline and Test sessions, we analyzed mean ERP amplitudes in predefined time windows based on the grand-average waveforms: P1 (90–140 ms), N1 (150–200 ms), P2 (220–270 ms), N2 (270–300 ms), and P300 (320–450 ms). Electrodes were selected a priori based on two criteria: (i) midline coverage and (ii) acceptable data quality (not noisy in more than 10 participants), and were further verified against the scalp topographies to ensure that the chosen sites captured the maximal component activity.

#### Pupil Analysis

Pupil data were preprocessed in R (v3.1.1) using RStudio (v0.98.1049). Baseline pupil diameter was defined for each trial as the mean pupil size during the 500 ms interval preceding stimulus-array onset. Pupil responses were then baseline-corrected by subtracting this value from each subsequent sample, yielding change in pupil diameter (mm) relative to baseline. Analyses were restricted to the Training sessions and conducted separately for stimulus-locked and feedback-locked responses. To assess early learning effects associated with reward-biased locations (20Lh, 50Lh, 50Hh, 80Hh), we averaged baseline-corrected pupil change in a 1000–1750 ms window after stimulus-array onset. To assess outcome-related effects, we averaged baseline-corrected pupil change in a 1000–1750 ms window after feedback onset, separately for high- versus low-reward trials.

## Acknowledgements

We thank all participants for their time and contribution to this study. We also thank Dr. Maja Dyhre Foldal for programming the experimental task, and Professor René Huster for valuable advice and discussions regarding ERP analysis.

## Author contributions

Conceptualization: OA, MS, TE; Methodology: OA, MS, TH, TE; Formal analysis: OA, MS, TH; Investigation: OA, MS, TH, TE; Resources: TE; Data curation: OA, MS, TH, TE; Project administration: OA, MS, TE; Funding acquisition: TE; Visualization: OA, MS, TH; Supervision: MS, TE; Writing-original draft: OA, MS; Writing-review and editing: OA, MS, TH, TE

## Supplementary Information

### Double target condition

Restricting the analysis to trials in which the two targets appeared specifically at the 80Hh and 20Lh locations and the participant successfully reported only one target resulted in very low trial counts. The table below reports, for each participant and session (Baseline, Test), the number of retained single-report trials in which the reported target was at 80Hh or 20Lh. After artifact rejection, the remaining trial numbers were generally insufficient to yield reliable averaged ERP waveforms. As a benchmark, at least ∼30 trials are typically recommended to obtain stable estimates even for relatively large components such as the P3 (Luck, 2005), and fewer trials are expected to be even more problematic for smaller components. Accordingly, we did not conduct or report inferential statistics on the N2 and P3 specifically for the strict 80Hh–20Lh pairing subset. Instead, we increased reliability by collapsing across single-report trials in which the reported target was at 80Hh versus 20Lh, irrespective of the location of the other (unreported) target.

| Subject_Num | Base_80_20_20 | Base_80_20_8<br>0 | Test_80_20_2<br>0 | Test_80_20_80 |
| --- | --- | --- | --- | --- |
| Subject101 | 26 | 8 | 21 | 15 |
| Subject102 | 4 | 15 | 6 | 8 |
| Subject103 | 6 | 18 | 1 | 6 |
| Subject104 | 13 | 16 | 12 | 12 |
| Subject105 | 29 | 14 | 19 | 19 |
| Subject106 | 10 | 26 | 7 | 30 |
| Subject108 | 18 | 9 | 18 | 3 |
| Subject110 | 12 | 16 | 20 | 5 |
| Subject111 | 1 | 14 | 2 | 11 |
| Subject112 | 3 | 18 | 1 | 16 |
| Subject114 | 15 | 10 | 8 | 6 |
| Subject115 | 20 | 15 | 23 | 13 |
| Subject118 | 19 | 5 | 18 | 10 |
| Subject121 | 3 | 24 | 5 | 15 |
| Subject122 | 25 | 9 | 22 | 7 |
| Subject123 | 30 | 7 | 30 | 6 |
| Subject125 | 11 | 30 | 5 | 15 |
| Subject126 | 13 | 18 | 13 | 10 |
| Subject127 | 10 | 3 | 3 | 1 |
| Subject128 | 21 | 15 | 18 | 14 |
| Subject129 | 8 | 11 | 8 | 13 |
| Subject131 | 27 | 7 | 27 | 7 |
| Subject132 | 11 | 26 | 11 | 10 |
| Subject133 | 20 | 20 | 16 | 16 |
| Subject203 | 4 | 30 | 2 | 14 |
| Subject205 | 6 | 19 | 3 | 14 |
| Subject206 | 10 | 10 | 7 | 11 |
| Subject207 | 12 | 24 | 10 | 23 |
| Subject210 | 17 | 13 | 14 | 11 |
| Subject211 | 22 | 12 | 17 | 16 |

